# Cocaine experience abolishes the motivation suppressing effect of CRF in the ventral midbrain

**DOI:** 10.1101/655936

**Authors:** Idaira Oliva, Melissa M. Donate, Merridee J. Lefner, Matthew J. Wanat

**Author notes:** **Corresponding Author:** Matthew J. Wanat, Neurosciences Institute, Department of Biology, University of Texas at San Antonio, One UTSA Circle, San Antonio, TX 78249.

## Abstract

Stress affects dopamine-dependent behaviors in part through the actions of corticotropin releasing factor (CRF) in the ventral tegmental area (VTA). For example, acute stress engages CRF signaling in the VTA to suppress the motivation to work for food rewards and promote drug seeking behavior. These diverging behavioral effects in food- and drug-based tasks could indicate that CRF modulates goal-directed actions in a reinforcer-specific manner. Alternatively, prior drug experience could functionally alter how CRF in the VTA regulates dopamine-dependent behavior. To address these possibilities, we examined how intra-VTA injections of CRF influenced cocaine intake and whether prior drug experience alters how CRF modulates the motivation for food rewards. Our results demonstrate that intra-VTA injections of CRF had no effect on drug intake when self-administering cocaine under a progressive ratio reinforcement schedule. We also found that a prior history of either contingent or non-contingent cocaine infusions abolished the capacity for CRF to reduce the motivation for food rewards. Furthermore, voltammetry recordings in the nucleus accumbens illustrate that CRF in the VTA had no effect on cocaine-evoked dopamine release. These results collectively illustrate that exposure to abused substances functionally alters how neuropeptides act within the VTA to influence motivated behavior.

## Introduction

Stress induces the release of corticotropin releasing factor (CRF), which is responsible for initiating the hormonal and physiological responses to stress [1]. CRF signaling within the mesocorticolimbic system also mediates the effects of stress on motivation and decision-making processes [2]. For example, CRF receptor activation within the ventral tegmental area (VTA) is required for the stress-induced reduction in the motivation to work for food rewards [3]. Acute stress also decreases the preference for high effort / large reward options in decision making tasks [4]. This alteration in effort-based decision making is blocked by antagonizing CRF receptors and recapitulated by administering CRF directly into the VTA [4]. Therefore, the actions of CRF in the ventral midbrain are critical for how stress regulates behavioral responding for natural rewards.

Whereas acute stress reduces the motivation to work for food rewards, stress promotes the reinstatement of drug seeking in a CRF-dependent manner [5]. Antagonizing CRF receptors in the VTA prevents stress-induced cocaine seeking [6,7]. Furthermore, intra-VTA injections of CRF are sufficient to reinstate cocaine seeking in the absence of stress [6,7], which together highlights the involvement of midbrain CRF signaling in how stress promotes drug-dependent behaviors.

How does CRF act within the VTA to suppress the motivation for food rewards and also enhance drug seeking behavior? One possibility is that CRF influences behavior in a reinforcer-specific manner. Since abused substances alter the electrophysiological effects of CRF on dopamine neurons [8-11], an alternative possibility is that prior drug experience changes how CRF in the VTA regulates dopamine-dependent behavior. Here, we performed experiments to determine whether the behavioral effects of CRF in the VTA are reinforcer-specific or regulated by prior drug intake. Rats were either (i) trained to nosepoke for cocaine infusions, (ii) received yoked cocaine infusions, or (iii) were drug naïve before they were trained to lever press for food rewards. We examined how intra-VTA CRF injections affected the motivation to work for cocaine infusions and food rewards under a progressive ratio (PR) reinforcement schedule. In this manner, we could determine if CRF differentially influenced the motivation to work for drug and food rewards within the same subjects. Additionally, we could ascertain if a prior history of contingent or non-contingent drug experience altered how CRF regulated the motivation for food rewards.

## Methods

### Subjects and surgery

All procedures were approved by the Institutional Animal Care and Use Committee at the University of Texas at San Antonio. Male Sprague-Dawley rats (Charles River, MA) were pair-housed upon arrival and given ad libitum access to water and chow and maintained on a 12-hour light/dark cycle. Surgeries were performed under isoflurane anesthesia on rats weighing between 300-350 g. For intracranial surgeries, rats were implanted with a bilateral guide cannula targeting the VTA (relative to bregma: 5.6 mm posterior; ± 0.5 mm lateral; 7.0 mm ventral). Rats used for voltammetry recordings were additionally implanted with carbon fiber electrodes in the nucleus accumbens (relative to bregma: 1.3 mm anterior; ± 1.3 mm lateral; 7.0 mm ventral) along with a Ag/AgCl reference electrode placed at a convenient location. All components were secured in place with cranioplastic cement. Rats were single housed following the intracranial implantation surgery and for the duration of the experiment. Intravenous jugular catheter surgeries were performed 1-3 weeks following the intracranial surgery. Rats assigned to the yoked cue control group underwent a mock catheter surgery in which they only received a backmount cannula implant. All animals were allowed to recover for at least 1 week following the catheter surgery before initiating training.

### Cocaine self-administration training

Cocaine self-administration sessions were performed in operant boxes (Med Associates) with grid floors, a houselight, two nosepoke ports with lights, and a tone generator, as described previously [12]. Behavioral sessions (1 hr) began with the illumination of the houselight and were performed only once per day. After completing the required number of nosepokes into the active port, rats received a 0.3 mg/kg i.v. infusion of cocaine, the tone and active nosepoke light turned on (5 s), which coincided with the initiation of a 20 s timeout period (houselight off). Nosepokes during the timeout or into the inactive port had no consequences. Rats were first trained to self-administer cocaine on a fixed ratio – 1 (FR1) reinforcement schedule. After rats completed at least 10 FR1 sessions and earned ≥ 10 infusions per day for at least two consecutive sessions, they were then trained to self-administer cocaine under a PR reinforcement schedule in which the operant requirement escalated according to the following equation: operant requirement = 5 * e^(infusion number * 0.2)^ – 5. Intra-VTA microinjections were performed once rats achieved < 15% variance in the infusions earned over three consecutive sessions. Rats failing to earn at least 6 infusions during PR sessions were excluded from the study. Yoked cocaine rats underwent 15 training sessions in which they received 23 infusions, which was based upon the performance of a rat that had self-administered cocaine. The yoked cocaine delivery was accompanied by the tone and active nosepoke light turning on (5 s) and the houselight turning off for 20s, which was identical to the conditions in rats that self-administered cocaine. Yoked cue rats underwent the identical training procedures as the yoked cocaine rats (including backmount attachment to the infusion tubing), though they did not receive cocaine infusions.

### Food lever press training

Lever press training for food rewards was performed after cocaine self-administration experiments, 15 sessions of non-contingent cocaine infusions (yoked cocaine), or 15 sessions of non-contingent cue presentations (yoked cue). Rats were placed and maintained on mild food restriction (∼15 g/day of standard lab chow) to target 90% free-feeding weight, allowing for an increase in weight of 1.5% per week. Experimental 45-mg food pellets (F0021, BioServ, NJ) were placed in their home cages on the day prior to the first training session to familiarize the rats with the food pellets. Behavioral sessions were performed in operant chambers that had grid floors, a house light, and contained a food tray and two cue lights above two retractable levers on a single wall. The cue lights and their corresponding levers were located on either side of the food tray. Rats were first trained to lever press for food rewards during sessions in which a single lever was presented throughout the duration of the session and a single lever press resulted in the delivery of the food reward. After rats successfully completed this training session (100 pellets earned within 90 mins), they were then trained on sessions in which both levers were extended and the cue light was illuminated over the active lever. Completion of the correct number of lever presses led to the delivery of the food reward, retraction of the levers, the cue and house lights turning off and a light over the food tray turning on for a 30 s inter-trial interval. Behavioral sessions consisted of a total of 60 trials and were performed only once per day. Rats were first trained to complete 60 trials on an FR1, FR2 and FR4 reinforcement schedule before progressing to a PR reinforcement schedule. PR sessions were identical to FR sessions except that the operant requirement on each trail (T) was the integer (rounded down) of 1.4^(T-1)^ lever presses starting at 1 lever press, as described previously [3,13]. Food PR sessions ended after 15 min elapsed without completion of the response requirement in a trial. Note that lever press responding is calculated as a rate to normalize behavioral performance across animals that differ in the duration of the food PR session. Intra-VTA microinjections were performed once rats achieved < 15% variance in the food pellets earned over three consecutive sessions.

### Voltammetry recordings

Chronically-implanted carbon-fiber microelectrodes were connected to a head-mounted voltammetric amplifier for dopamine detection in behaving rats using fast-scan cyclic voltammetry as described previously [3,12,14]. The potential applied to the carbon fiber was ramped in a triangle wave from −0.4 V (vs Ag/AgCl) to +1.3 V and back at a rate of 400 V/s during a voltammetric scan and held at −0.4 V between scans at a frequency of 10 Hz. Dopamine was isolated from the voltammetry signal using chemometric analysis [15], using a standard training set accounting for dopamine, pH, and background drift. The dopamine concentration was estimated based on the average post-implantation sensitivity of electrodes (34 nA/μM) [14]. Peak cocaine-evoked dopamine release was calculated for the 100 s following a single 1.8 mg/kg i.v. cocaine infusion [16].

### Intra-VTA microinjections and data analysis

Rats received bilateral 0.5 μl injections of CRF (Bachem) or ACSF (Tocris) into the VTA at a rate of 250 nl/min. The injector was removed from the brain after at least 1 min had elapsed since the end of the infusion. Rats were placed in their homecage for 20 min before starting behavioral sessions. The dose-dependent effect of CRF injections on behavior were assessed in a counterbalanced manner. Intra-VTA injections were separated by at least one session in which no injections were performed. Additionally, subsequent injections were not performed until the rewards earned during the behavior-only PR sessions came within 15% of the average rewards earned during the three baseline sessions. Statistical analyses utilized a one-way ANOVA followed by post-hoc Dunnett’s test relative to the ACSF injection. The Geisser-Greenhouse correction was applied to address unequal variances between the treatments for repeated measures ANOVAs.

### Histology

Histology was performed to verify the placement of guide cannula and voltammetry electrodes (Supplemental Fig. 1). Rats were intracardially perfused with 4% paraformaldehyde and brains were removed and post-fixed in the paraformaldehyde solution for at least 24 h. Brains were subsequently placed in 15 % and 30% sucrose solutions in phosphate-buffered saline. Brains were then flash frozen in dry ice, coronally sectioned and stained with cresyl violet.

## Results

Male rats were trained to nosepoke for cocaine infusions and then to lever press for food rewards. In this manner we could ascertain how CRF acts within the VTA to influence the motivation to work for drug and food rewards in the same animals (Fig. 1A). Rats self-administered cocaine (0.3 mg/kg i.v.) on an FR1 reinforcement schedule (mean 11.2 ± 0.7 training sessions, n = 10 rats) before self-administering cocaine on a PR reinforcement schedule (mean 10.1 ± 1.1 training sessions). Animals then received bilateral intra-VTA injections of CRF (0 −1 µg) prior to cocaine PR sessions. Local injections of CRF into the VTA had no effect on the number of cocaine infusions earned (one-way repeated measures ANOVA F_(2.3,20.4)_ = 0.2, p = 0.83; Fig. 1B), the number of active nosepokes (one-way repeated measures ANOVA F_(2.1,18.8)_ = 0.1, p = 0.88; Fig. 1C), or the number of inactive nosepokes during cocaine PR sessions (one-way ANOVA F_(1.4,19.9)_ = 0.3, p = 0.70; Fig. 1D). Rats were then trained to lever press for food rewards, before assessing how intra-VTA CRF injections (0 −1 µg) affected the motivation to work for food rewards on a PR reinforcement schedule (Fig. 1A). Our results demonstrate that CRF did not influence the number of food rewards earned (one-way repeated measures ANOVA F_(2.0,17.8)_ = 0.6, p = 0.55; Fig. 1E), the rate of active lever presses (one-way repeated measures ANOVA F_(2.4,21.9)_ = 0.4, p = 0.71; Fig. 1F), or the rate of inactive lever presses (one-way repeated measures ANOVA F_(2.1,18.7)_ = 1.1, p = 0.37; Fig. 1G). Although CRF acts within the VTA to inhibit the motivation to work for food rewards in drug-naïve animals [3], we find that CRF does not affect the motivation to work for cocaine infusions or for food rewards in animals with prior drug experience.

**Figure 1.**
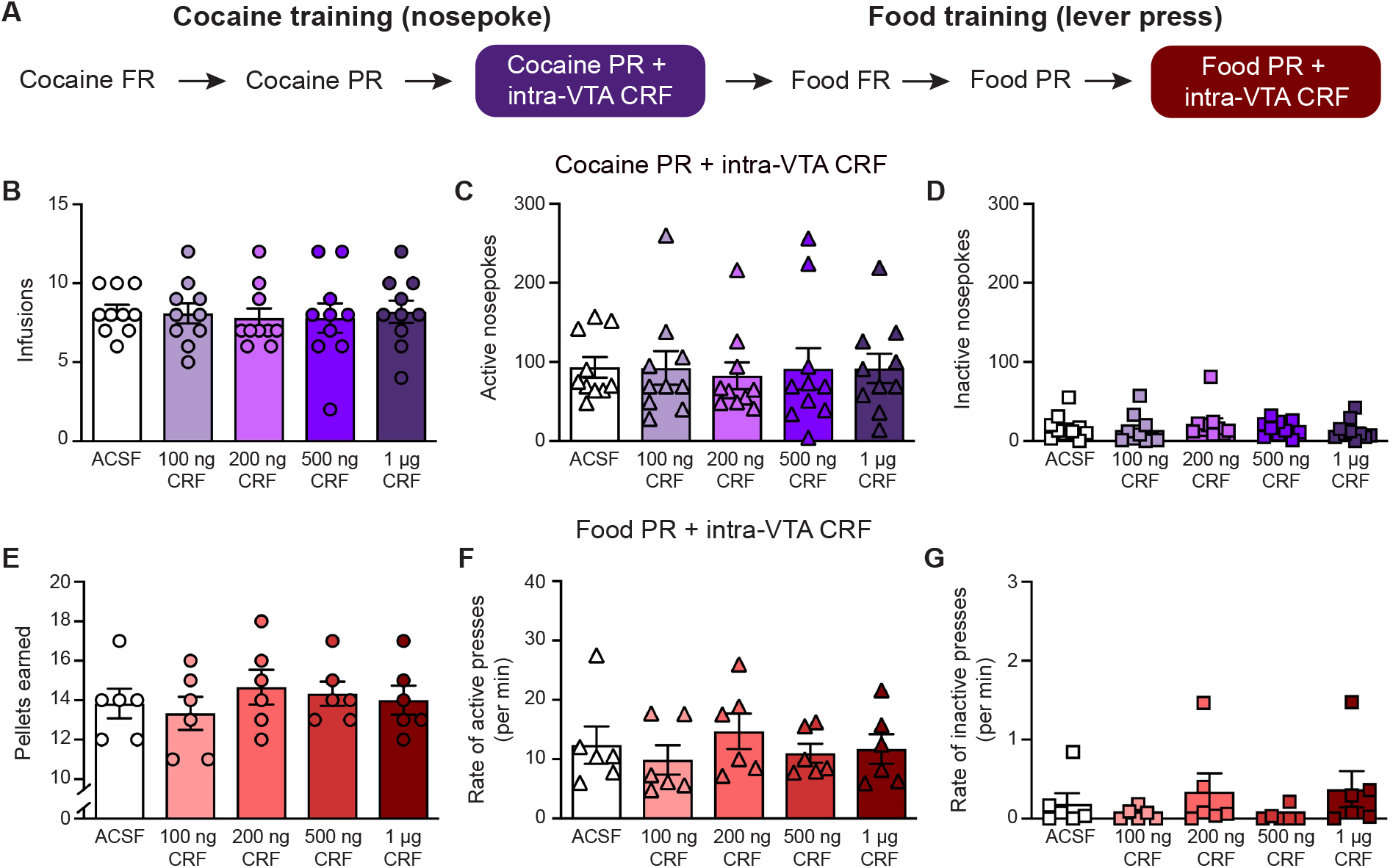
CRF does not affect the motivation for drug or food rewards following cocaine self-administration. (A) Outline of training for rats self-administered cocaine. (B-D) Effect of intra-VTA injections of CRF on cocaine infusions (B), active nosepokes (C), and inactive nosepokes (D) during cocaine self-administration sessions under a PR reinforcement schedule. (E-G) Effect of intra-VTA injections of CRF on food pellets earned (E), rate of active lever presses (F), and rate of inactive lever presses (G) during operant responding for food rewards under a PR reinforcement schedule.

The inability for CRF to regulate the motivation for food rewards following a history of cocaine self-administration could be due to the pharmacological actions of cocaine or alternatively could require contingent drug intake. To address these possibilities, a separate cohort of rats received yoked cocaine infusions along with the corresponding cocaine delivery cues (n = 10 rats). Following 15 sessions of non-contingent cocaine infusions, rats were trained to lever press for food pellets before we assessed how intra-VTA CRF injections influenced the motivation to work for food rewards (Fig. 2A). Identical to the results from the cocaine self-administering rats (Fig. 1), CRF injections had no effect on the number of food pellets earned (one-way repeated measures ANOVA F_(2.1,19.1)_ = 2.0, p = 0.16; Fig. 2B), the rate of active lever presses (one-way repeated measures ANOVA F_(2.8,25.4)_ = 3.0, p = 0.05; Fig. 2C), or the rate of inactive lever presses during food PR sessions in rats that had received yoked cocaine infusions (one-way repeated measures ANOVA F_(1.9,17.0)_ = 0.6, p = 0.58; Fig. 2D). Therefore, the pharmacological effects of cocaine are sufficient to prevent the avolitional influence of CRF in the VTA.

**Figure 2.**
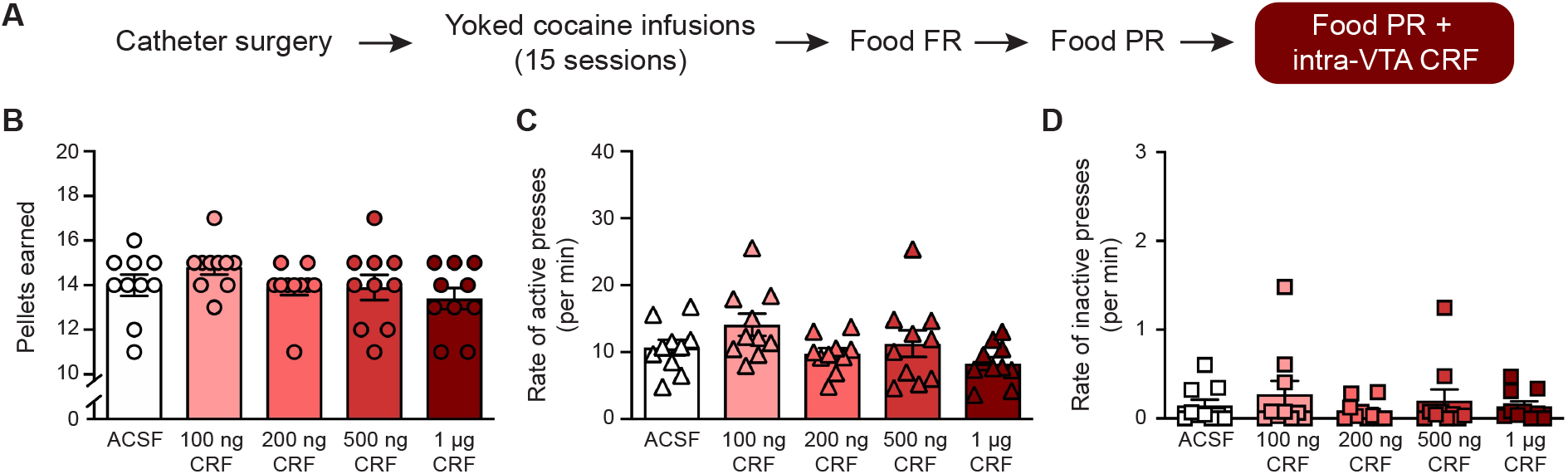
CRF does not affect the motivation for food rewards following non-contingent cocaine infusions. (A) Outline of training for yoked cocaine treated rats. (B-D) Effect of intra-VTA injections of CRF on food pellets earned (B), the rate of active lever presses (C), and the rate of inactive lever presses

We next sought to exclude the possibility that the catheter surgery and/or exposure to the cues contributed to the change in the capacity for CRF to regulate motivation. To address this, a third group of rats underwent a mock catheter surgery followed by sessions in which they received the yoked presentation of cues (n = 12 rats). The yoked cue presentations were identical to the cues used from the self-administration / yoked cocaine experiments, except that no drug was delivered. Following 15 yoked cue presentation sessions, rats were trained to lever press for food rewards before assessing how intra-VTA CRF injections influenced the motivation to work for food (Fig. 3A). We found that CRF dose-dependently reduced the number of food pellets earned (one-way repeated measures ANOVA F_(2.5,27.0)_ = 13.0, p < 0.0001; post-hoc Dunnett’s test relative to ACSF: 100 ng, q_11_ = 3.0, p < 0.05; 200 ng, q_11_ = 3.5, p < 0.05; 500 ng q_11_ = 4.6, p < 0.01; 1 µg q_11_ = 4.4, p < 0.01; Fig. 3B). The suppression in rewards earned was accompanied by a dose-dependent reduction in the rate of active lever presses (one-way repeated measures ANOVA F_(2.2,23.9)_ = 7.8, p = 0.0019; post-hoc Dunnett’s test relative to ACSF: 100 ng, q_11_ = 2.4, p > 0.05; 200 ng, q_11_ = 2.5, p > 0.05; 500 ng q_11_ = 4.1, p < 0.01; 1 µg q_11_ = 3.4, p < 0.05; Fig. 3C). There was a main effect of intra-VTA CRF injections on the rate of inactive lever presses, though post-hoc analyses found no difference between ACSF injections and CRF injections at any of the tested doses (one-way repeated measures ANOVA F_(1.8,20.0)_ = 3.9, p = 0.04; Fig. 3D).

**Figure 3.**
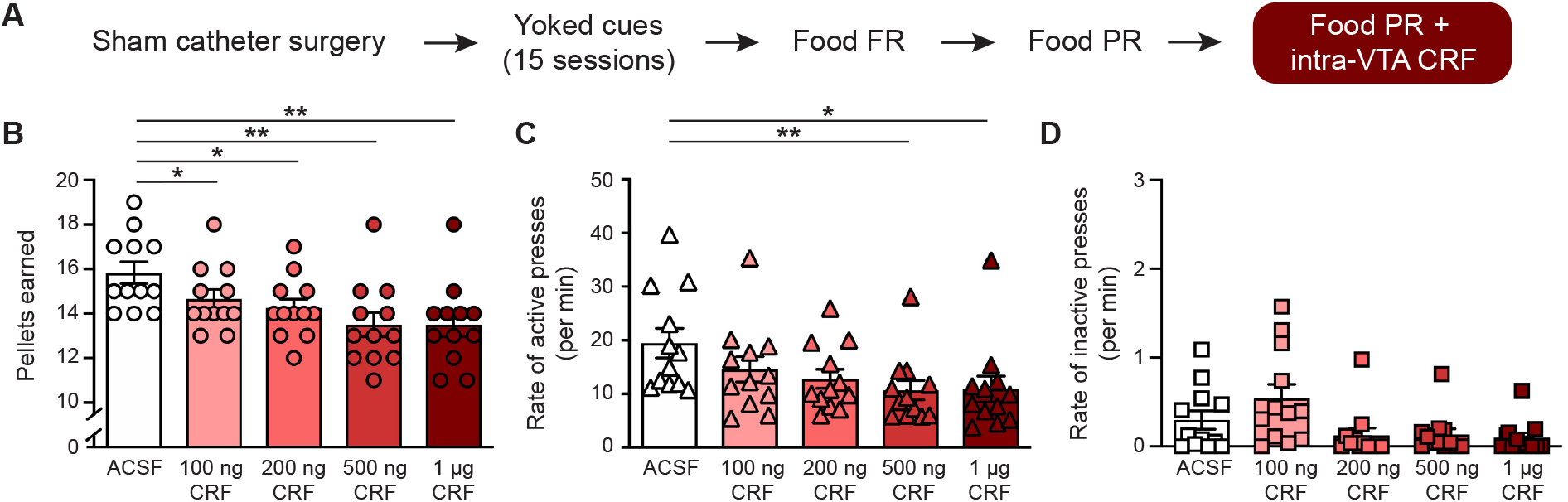
CRF inhibits the motivation to work for food rewards in drug-naïve animals. (A) (A) Outline of training for yoked cue rats. (B-D) Effect of intra-VTA injections of CRF on food pellets earned (B), rate of active lever presses (C), and rate of inactive lever presses (D) during operant responding for food rewards under a PR reinforcement schedule in rats that had received yoked presentations of cues. * *p* < 0.05, ** *p* < 0.01, *** *p* < 0.001.

The inability for CRF to affect the motivation for food rewards in cocaine-treated rats was not due to intrinsic differences in behavioral responding between the groups. Specifically, there was no difference between the yoked cue, yoked cocaine, and cocaine self-administration groups in the rewards earned (one-way ANOVA F_(2,29)_ = 1.0, p = 0.38; Fig. 4A), the rate of active presses (one-way ANOVA F_(2,29)_ = 1.0, p = 0.40; Fig. 4B), or the rate of inactive lever presses (one-way ANOVA F_(2,29)_ = 0.2, p = 0.81; Fig. 4C) during the baseline PR sessions prior to intra-VTA CRF injections. Together, these data demonstrate that CRF acts within the midbrain to inhibit the motivation to work for food rewards only in drug-naïve animals.

**Figure 4.**
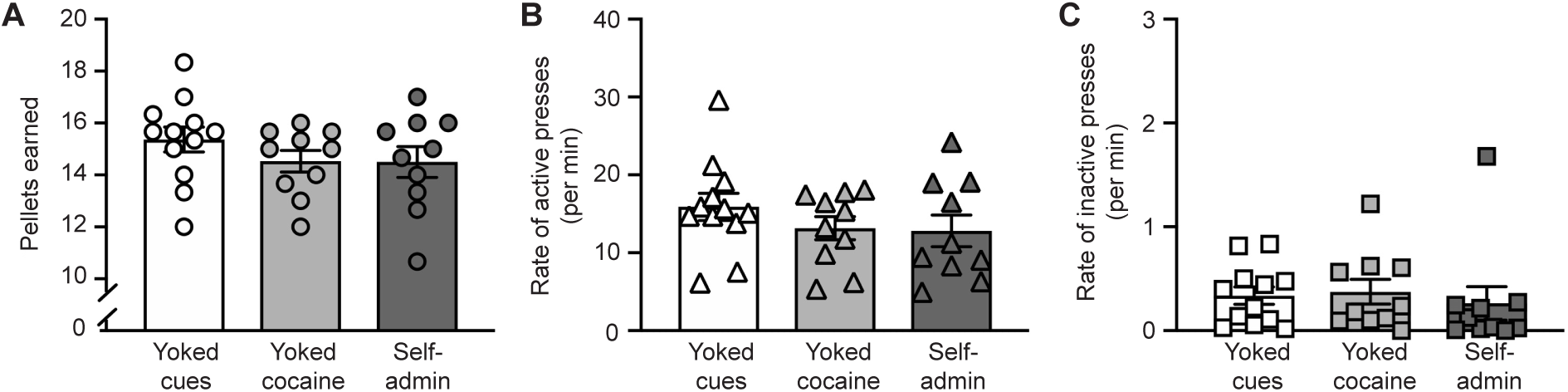
No behavioral differences between groups during baseline food PR sessions. (A-C) The relative behavioral performance between the cocaine self-administration, yoked cocaine, and yoked cue rats during baseline PR sessions. There was no difference in food pellets earned (A), the rate of active lever presses (B), and rate of inactive lever presses (C).

Dopamine transmission in the nucleus accumbens is required for engaging in high-effort behaviors [17]. Conversely, suppressing dopamine neuron activity at the time of the reward delivery reduces the motivation to work for rewards [18]. In drug naïve animals, CRF acts within the VTA to attenuate dopamine release to food rewards [3]. Since midbrain CRF does not influence motivation following exposure to cocaine, we anticipated that CRF would not affect dopamine release to drug rewards in cocaine experienced animals. To test this prediction, we performed voltammetry recordings in the nucleus accumbens of dopamine release to an infusion of cocaine (1.8 mg/kg i.v.). Rats were habituated to this procedure (minimum 10 prior cocaine infusions) before assessing how intra-VTA injections of CRF influenced cocaine-evoked dopamine release. Although dopamine release to drug rewards is controlled by neuronal activity within the VTA [16,19], CRF in the VTA had no effect on dopamine release to cocaine infusions (one-way ANOVA F_(4,22)_ = 0.4, p = 0.84; n = 6 electrodes, Fig. 5A-C). These findings along with our prior work [3], illustrates the capacity for midbrain CRF to regulate reward-evoked dopamine release is related to its ability to influence motivated behavior.

**Figure 5.**
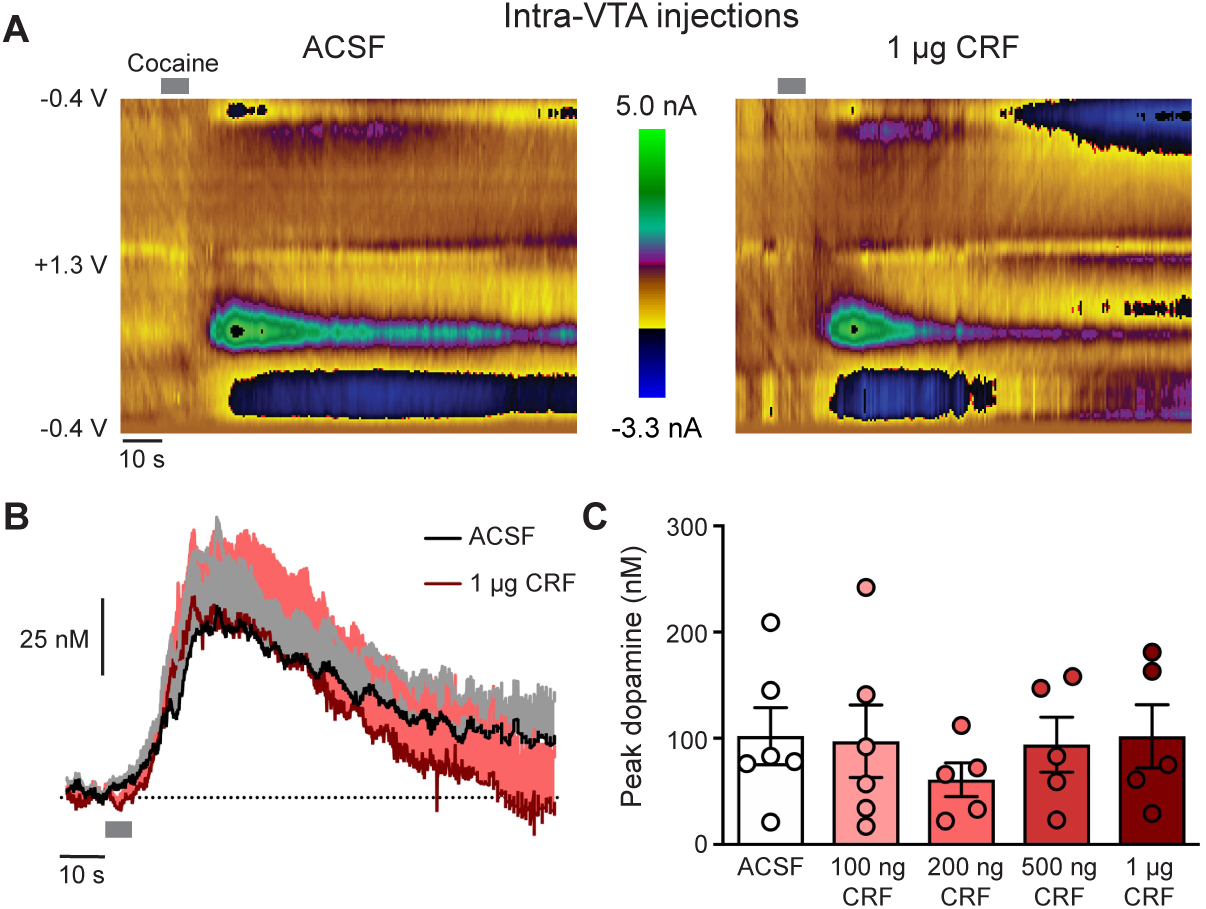
Intra-VTA injections of CRF do not affect cocaine-evoked dopamine release. (A) Representative colorplots of voltammetry recordings from a single electrode of cocaine-evoked dopamine release after intra-VTA injections of ACSF (left) or 1 µg CRF (right). (B) Average dopamine response to the cocaine infusion across electrodes (C) Peak dopamine release to the cocaine infusion.

## Discussion

Acute stress influences a host of behaviors through the actions of CRF in the VTA, with effects on motivation, decision-making, subsequent drug intake, and drug seeking [3,4,6,7,20-22]. Intriguingly, midbrain CRF suppresses dopamine-dependent behaviors in drug-naïve animals [3,4], whereas CRF promotes dopamine-dependent behaviors in drug-experienced animals [6,7,20-22]. These conflicting reports indicate that the behavioral effects of CRF in the midbrain are either reinforcer-specific, or alternatively depend upon prior drug experience. Our results support the latter, as we demonstrate that the capacity for midbrain CRF to reduce the motivation for food rewards is lost following a history of cocaine self-administration or non-contingent cocaine infusions. Although CRF promotes the reinstatement of drug seeking in rats that have undergone extinction training [6,7], we find that midbrain CRF does not affect the motivation to work for cocaine during self-administration sessions. Therefore, the functional impact of CRF in the VTA on drug-dependent behaviors may depend upon if the reinforcer can be earned.

The motivation to work for rewards is enhanced by augmenting mesolimbic dopamine transmission and attenuated by inhibiting dopamine signaling [23-26]. In particular, motivation in appetitive tasks is influenced by dopamine transmission at the time of the reward delivery [18]. Large rewards evoke greater dopamine release relative to small rewards, which parallels the greater effort rats will exert to obtain larger rewards [3]. Conversely, the CRF-mediated reduction in motivation to work for food rewards is accompanied by a decrease in reward-evoked dopamine release [3]. We found that intra-VTA injections of CRF do not affect the motivation to work for cocaine infusions or dopamine release to cocaine infusions. Together, these findings demonstrate that the motivation to work for a given reward is intimately linked to how the dopamine system responds to the reward delivery.

Although midbrain CRF attenuates reward-evoked dopamine release in drug-naïve animals [3], it is likely that CRF is not solely acting upon dopamine neurons within the VTA. Electrophysiological studies have identified numerous cell-autonomous effects of CRF on dopamine neurons, including changes in firing rate [27], excitatory currents [10,28,29], and inhibitory currents [8]. CRF also modulates the firing rate of VTA GABA neurons [30] and regulates presynaptic inhibitory and excitatory input onto dopamine neurons [11,31]. These electrophysiological effects are mediated by activation of the CRF-R1 [8,27,31] and CRF-R2 [10,29] or both [11]. Recent evidence illustrates regional differences in how CRF is released within the VTA [22] and that CRF regulates afferent input in a pathway-selective manner [3]. Together, these prior studies indicate that the behavioral effects of CRF in the VTA likely arises from a complex interplay of the neuropeptide’s actions on a diverse set of targets within the midbrain.

Exposure to abused substances elicits an array of intrinsic and synaptic changes within the VTA [32]. Prior drug experience increases CRF receptor levels within the VTA [9,33,34], and alters the electrophysiological effects of CRF on VTA dopamine neurons [8,10,11,31]. As such, the cocaine-mediated loss in the capacity for CRF to regulate motivation could be mediated by a drug-induced alteration in which CRF engages a local circuit to reverse its inhibitory influence on reward-evoked dopamine release. Collectively, our data illustrates that both contingent and non-contingent exposure to abused drugs functionally alters the manner by which CRF acts within the midbrain to control dopamine-dependent behavior.

## Acknowledgements

I.O., M.M.D., and M.J.F. performed the experiments. I.O., M.M.D., M.J.F., and M.J.W. analyzed the data. I.O. and M.J.W. designed the experiments. M.J.W wrote the first draft of the manuscript and I.O., M.M.D, and M.J.F. made revisions. This work was funded by NIH grants to M.J.W. (DA033386 and DA042362).

