## Supplementary Figure 1 for "Cocaine experience abolishes the motivation suppressing effect of CRF in the ventral midbrain"

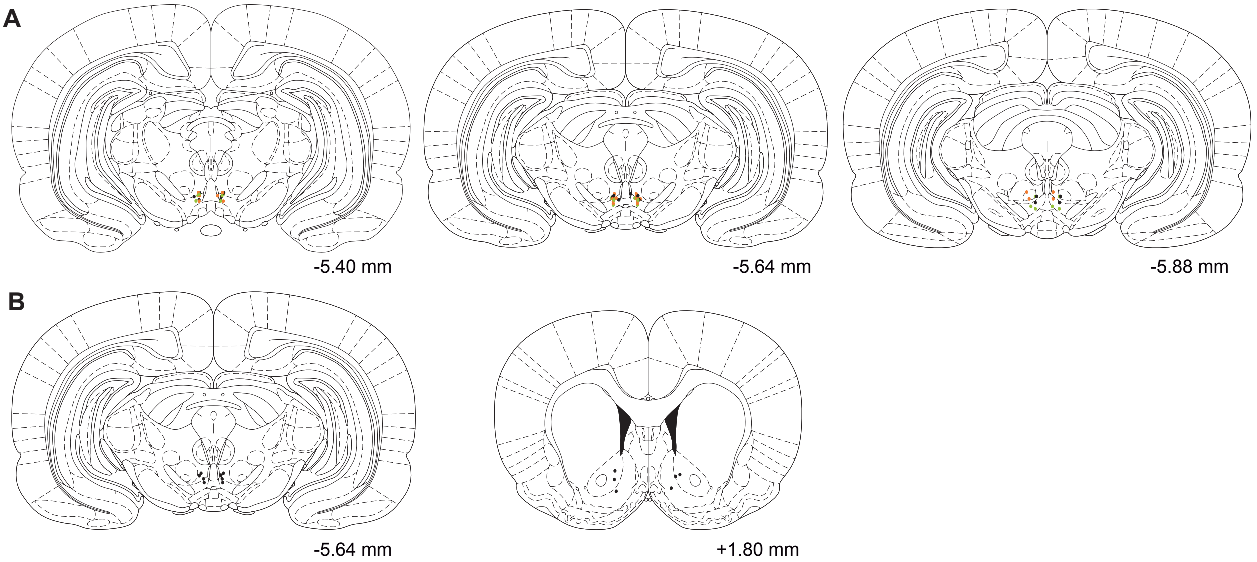


**Supplementary Figure 1. Histology.** (A) Placement of guide cannulas for rats that had self-administered cocaine (black), received yoked cocaine infusions (green), or received yoked cues (orange). (B) Placement of guide cannulas and voltammetry electrodes.
